# Temporal Changes in Innate and Adaptive Immunity During Sepsis as Determined by ELISpot

**DOI:** 10.1101/2023.12.14.571668

**Authors:** J Unsinger, D Osborne, AH Walton, E Han, L Sheets, MB Mazer, KE Remy, TS Griffith, M Rao, VP Badovinac, SC Brackenridge, I Turnbull, Philip A Efron, LL Moldawer, CC Caldwell, RS Hotchkiss

**Affiliations:** Department of Anesthesiology, Washington University School of Medicine, St. Louis, Missouri 63110 USA; Department of Pediatrics, Case Western Reserve University School of Medicine, Cleveland, Ohio 44106USA; Department of Urology, University of Minnesota Medical 47 School, Minneapolis, Minnesota 55455, USA; Center for Immunology, University of Minnesota Medical School, Minneapolis, Minnesota 55455 USA; Minneapolis VA Healthcare System, Minneapolis, Minnesota 55417, USA; Department of Pediatrics, University of Iowa Carver College of Medicine; Department of Pathology, University of Iowa Carver College of Medicine, Iowa City, Iowa 52242 USA; Experimental Pathology Ph.D. Program, University of Iowa Carver College of Medicine, Iowa City, Iowa 52242 USA; Department of Surgery, Harborview Medical Center, University of Washington School of Medicine, Seattle, Washington 98133 USA; Department of Surgery, Washington University School of Medicine, St. Louis, Missouri 63110 USA; Sepsis and Critical Illness Research Center, Department of Surgery, University of Florida College of Medicine, Gainesville, Florida 32610 USA; Department of Surgery, University of Cincinnati College of Medicine, Cincinnati, Ohio 45267, USA

**Keywords:** innate immunity, adaptive immunity, IL-7, arginine, corticosteroids

## Abstract

**Background:** The inability to evaluate host immunity in a rapid quantitative manner in patients with sepsis has severely hampered development of novel immune therapies. The ELISpot assay is a *functional* bioassay that measures the number of cytokine-secreting cells and the relative amount of cytokine produced at the single-cell level. A key advantage of ELISpot is its excellent dynamic range enabling a more precise quantifiable assessment of host immunity. Herein, we tested the hypothesis on whether the ELISpot assay can detect dynamic changes in both innate and adaptive immunity as they often occur during sepsis. We also tested whether ELISpot could detect the effect of immune drug therapies to modulate innate and adaptive immunity.

**Methods:** Mice were made septic using sublethal cecal ligation and puncture (CLP). Blood and spleens were harvested serially and *ex vivo* IFN-γ and TNF-α production were compared by ELISpot and ELISA. The capability of ELISpot to detect changes in innate and adaptive immunity due to *in vivo* immune therapy with dexamethasone, IL-7, and arginine was also evaluated.

**Results:** ELISpot confirmed a decreased innate and adaptive immunity responsiveness during sepsis progression. More importantly, ELISpot was also able to detect changes in adaptive and innate immunity in response to immune-modulatory reagents, for example dexamethasone, arginine, and IL-7 in a readily quantifiable manner, as predicted by the reagents known mechanisms of action. ELISpot and ELISA results tended to parallel one another although some differences were noted.

**Conclusion:** ELISpot offers a unique capability to assess the functional status of both adaptive and innate immunity over time. The results presented herein demonstrate that ELISpot can also be used to detect and follow the *in vivo* effects of drugs to ameliorate sepsis-induced immune dysfunction. This capability would be a major advance in guiding new immune therapies in sepsis.

## INTRODUCTION

Sepsis, a life-threatening organ dysfunction caused by a dysregulated host response to infection, is the most common cause of mortality in intensive care units in the United States (1). Both pro- and anti-inflammatory processes are initiated simultaneously in sepsis and their intensity and duration vary throughout the course of the disease (2-4). The lack of an effective means to evaluate the intensity of the host immune status in sepsis in a rapid quantitative method has severely hampered the development of novel immune therapies that could potentially improve outcomes. This absence of a method to immune endotype septic patients is also undoubtedly a contributing factor in the failure of numerous drug therapy trials in sepsis. For example, administration of corticosteroids to patients in refractory septic shock is likely to be beneficial in the subset of patients who are in the more hyper-inflammatory phase but may well be detrimental to patients who are in the more immunosuppressive phase of the disorder (5,6). Although there have been considerable efforts to develop methods to immune endotype septic patients (e.g., using transcriptomics), these methods often do not reflect the *functional state* of host immunity, lack dynamic range, and are more indicative of molecular processes rather than host immunity. Additionally, the data may not be available in a clinically relevant timeframe. We also believe that the relative balance between pro- and anti-inflammatory processes is dynamic over the course of illness, so methods that only allow sampling at a single time point may be of limited utility in guiding therapeutics.

Recently, investigators have argued that monitoring the immune status of septic patients is a *necessary precondition* for administration of the various immune therapies (e.g., corticosteroids, GM-CSF, IFN-γ, anti-programmed cell death 1, and IL-7), that are being used and/or tested in sepsis. The ELISpot assay is a *functional* bioassay that measures the number of cytokine-secreting cells at the single-cell level (9). A key advantage of ELISpot is that the assay has excellent dynamic range. ELISpot can detect as little as one cytokine secreting cell in 100,000 cells or as many as several thousand cytokine secreting cells per individual chamber well. Since the size of each spot represents the amount of cytokine made by an individual cell, it is also possible to compare relative cytokine production on a per cell basis by determining the area of each spot. Importantly, ELISpot is technically easier to perform than several other methods used to quantitate cytokine production such as flow cytometry. Additionally, the ELISpot requires less blood than flow cytometry–based methods of assessing immune function, making it more amenable to multiple timepoint sampling or where blood sampling volume is limited, such as with pediatric patients. Furthermore, because ELISpot can quantitate the relative amount of cell cytokine production over an extended period (20-22 hours) it is more sensitive at detecting differences in the study groups compared to other methods employing shorter incubation times.

Another major advantage of ELISpot is that it can independently assess the functional status of the two major arms of immunity (10,11). For example, the functional state of *adaptive* immunity can be evaluated by anti-CD3/CD28 mAb stimulated IFN-γ production while *innate* immunity can be assessed by lipopolysaccharide (LPS) stimulated monocyte production of TNF-α. An important and potentially clinically relevant advantage of ELISpot is its ability to examine the effect of different immune active drug therapies to modulate the patients’ immune response *in vitro* using diluted whole blood sample which closely mimics the septic milieu. Therefore, ELISpot might be useful in selecting drug therapies that are more likely to be efficacious *in vivo* in septic patients because of their beneficial effects in the *in vitro* assay.

The purpose of this study was to characterize the temporal evolution of the adaptive and innate immune response to sepsis from the early initial phase to the later resolution phase of sepsis in a clinically relevant, mouse cecal ligation and puncture (CLP) mouse model of sepsis. Although the mouse immune system has many differences compared to the human immune system, the mouse CLP model of sepsis does exhibit a number of changes that are similar to human sepsis including activating both hyper- and hypo-inflammatory changes in innate and adaptive immunity (12-15). Secondly, we wished to determine if the ELISpot assay could detect the effect of *in vivo* administration of immune modulatory drug therapies to alter the sepsis immune response. Development of a clinically applicable test that could detect the effect of immune modulatory agents on the functional state of innate and adaptive immunity would be a potentially major advance. Many immune cells become activated during sepsis and upregulate the expression of adhesion molecules that facilitate the migration of the cells from the circulation and localize at sites of infection and inflammation. Therefore, examination of circulating immune cells may fail to adequately reflect the host immune response occurring in many organs. To better characterize the immune response in circulating versus tissue-based immune cells, the ELISpot assay was performed using either mouse whole blood or splenocytes.

## MATERIALS AND METHODS

Eight- to ten-week-old male C57BL/6 mice purchased from The Jackson Laboratory (Bar Harbor, ME) were employed for all studies. Male mice were used to avoid potential changes in immune response that may occur during the estrous cycle. Intra-abdominal peritonitis was induced with the cecal ligation and puncture (CLP) model, as described previously (16,17). Briefly, mice were anesthetized with isoflurane, and a midline abdominal incision was performed. The cecum was identified, ligated, and punctured once with a 27-g needle. The level of injury to the cecum (the amount of cecum ligated) was directed at inducing a mild to moderate degree of peritonitis and sepsis (∼LD_10_ at 72 hours) with about 1/3 of the cecum ligated and punctured. The only exception to this protocol was in the study examining the effect of sepsis at the early time point of 6 hours (**Fig. 1**). Here, one half of the cecum was ligated and punctured (∼LD_80_ at 72 hours). The abdomen was closed in two layers, and 1 ml of 0.9% saline as well as buprenorphine ER (0.01 mg/kg body weight) for pain relief were administered subcutaneously. Naive mice which did not undergo surgery were used as controls. All animal procedures were approved by the Washington University School of Medicine Animal Studies Committee.

**Figure 1.**
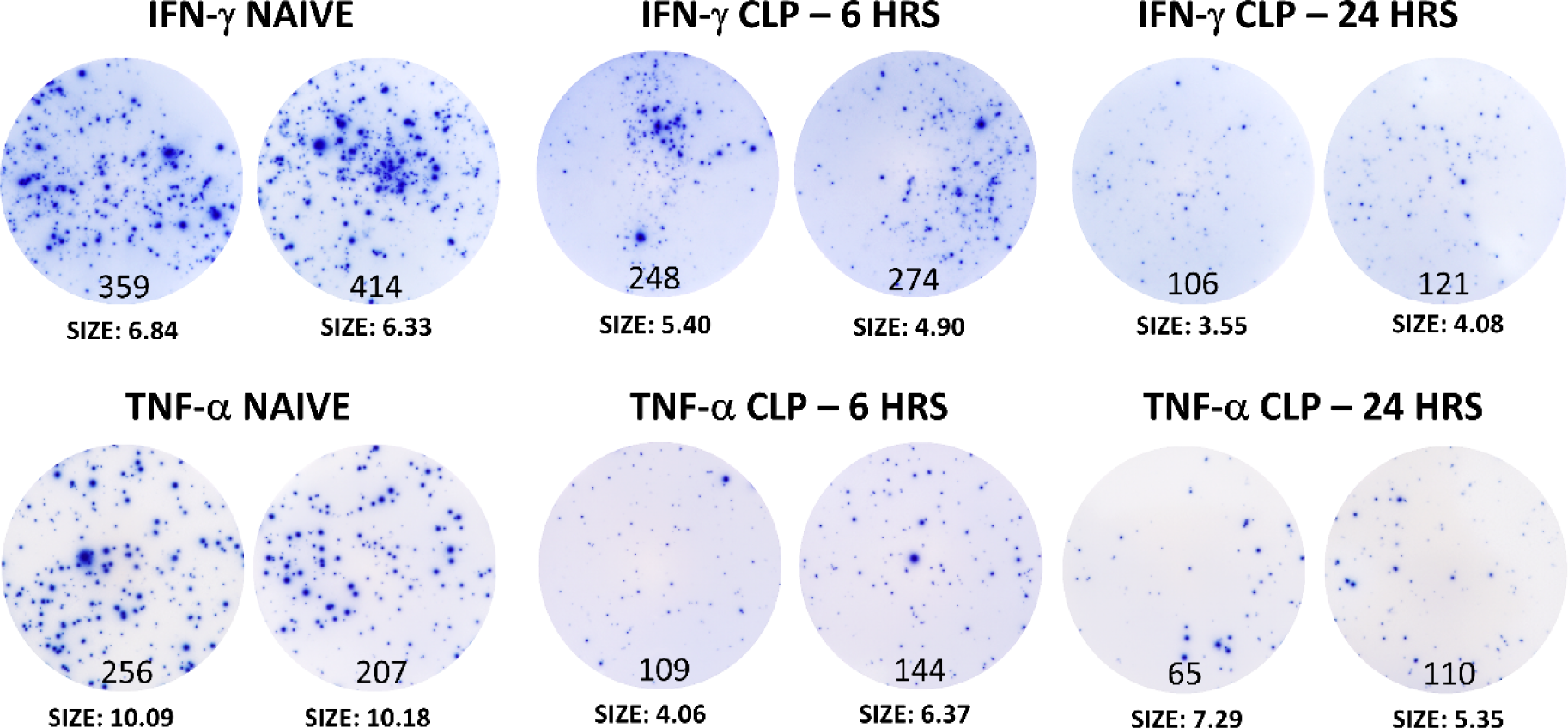
Rapid onset of innate and adaptive immune suppression in septic splenocytes. Mice underwent CLP and splenocytes were harvested 6 and 24 hours after induction of sepsis. Equal numbers of splenocytes from control mice and septic mice were stimulated with anti-CD3/anti-CD28 mAb for IFN-γ or LPS for TNF-α ELISpot assay. Each spot represents an immune cell producing either IFN-γ or TNF-α. The number of spots is depicted at the bottom of each circle and shows the marked loss in the number of cytokine-producing cells after CLP. There is also a decrease in the spot size representing a decrease in the amount of cytokine produced on a per cell basis. Tests were run in duplicate. Results are representative of 6 naïve mice, 6 mice at six hours, and 3 mice at 24 hours. The CLP surgery in these mice was more severe and involved tying off approximately one half of the cecum as opposed to one third of the cecum for all other CLP studies. The more severe CLP model was used to determine how quickly a more fulminant sepsis model would be detectable.

### Flow cytometric quantitation of absolute cell counts

Total cell counts per spleen were determined via the Vi-Cell counter (Beckman Coulter, Fullerton, CA) (16). The percentages of individual cell phenotypes (e.g., CD4, CD8, B cells, Monocytes/Macrophages and dendritic cells) were determined via flow cytometric analysis as previously described (16,17). Determination of cell populations were based on location by FSC and SSC as well as a combination of individual cell markers: CD4 T cells (CD3^+^, CD4^+^), CD8 T cells (CD3^+^, CD8^+^), B cells (B220^+^), Monocytes/Macrophages (B220^-^, CD11b^+^, CD11c^-^, Ly6G^-^), Dendritic cells (B220^-^, CD11b^+^, CD11c^+^, Ly6G^-^). Thus, the absolute cell counts for each splenic subset population were calculated, as described previously (16,17).

### Quantitation of IFN-γ and TNF-α production via ELISA

ELISA was performed to determine the impact of various immune treatments on the quantity of cytokines produced by T cells and monocytes/macrophages upon *ex vivo* stimulation as described previously (18,19). 5×10^6^ splenocytes were added to each well of a 24-well plate and stimulated with anti-CD3 and -CD28 mAb (1μg/ml and 5μg/ml, respectively; Biolegend) for IFN-γ production or LPS (1μg/ml, Enzo, Farmingdale, NY) for TNF-α production in 1 ml of complete RPMI1640 (10% FCS, Glutamine, non-essential amino acids, Pen/Strep) overnight. The supernatants were obtained and IFN-γ and TNF-α concentrations were quantified using ELISA MAX Standard Sets (Biolegend; cat.# 430801 for IFN-γ, cat.#430901 for TNF-α) employing the μQuant Scanning Microplate Spectrophotometer (Bio-Tek Instruments, Winooski, VT) as described (18,19)

### Quantitation of IFN-γ and TNF-α production via ELISpot

ELISpot assay was performed using either whole blood or splenocytes as previously reported (18,19). For the whole blood ELISpot, blood was collected by cardiac puncture in the presence of heparin. The whole blood was diluted 1:10 using CTL-test medium (Cellular Technology Ltd; Cleveland, OH) augmented with 2 mM glutamine. 50 μl of the diluted blood was added to a well containing 150 μl of the same media. For the splenocyte ELISpot, a different cell number/well was used when assaying for IFN-γ (2.5×10^4^ cells/well) or TNF-α (15×10^3^ cells/well). Cells were plated in duplicate in complete RPMI. For both the whole blood and splenocyte assays, cells were stimulated overnight with anti-CD3/CD28 mAb (1μg/ml and 5μg/ml, respectively) for IFN-γ production or LPS (2.5 ng/ml) to stimulate the production of TNF-α. Mouse ELISpot kits from Cellular Technology Ltd. (Cleveland, OH) were used following the manufacturer’s instructions. Following development, images were captured and analyzed on an ImmunoSpot S6 Universal-V Analyzer (Cellular Technologies Ltd).

### Modulation of the septic immune response with dexamethasone, IL-7, and arginine

Septic mice were treated with immune modulatory drugs with known activity on innate and adaptive immune cells (20-23). Dosage of drugs was as follows: IL-7 (2.5 μg in saline subcutaneously, per dose; kindly gifted by Dr. Michel Morre, RevImmune), dexamethasone (50 μg in saline subcutaneously, per dose; Hikma pharmaceuticals - purchased from Barnes Jewish Hospital pharmacy), L-arginine (2.5 mg intraperitoneally in saline per dose; Sigma-Aldrich, St. Louis, MO, Cat. No. A8094, lot no. SLBN0906V). Experimental details on the timing of administration of these three drugs are provided in the results section and respective figure legends.

### Statistical analysis

Data were analyzed with the statistical software Prism (GraphPad, San Diego, CA, USA). Data are reported as the mean ± SEM. For comparison of two groups, the Student’s *t*-test was employed. One-way ANOVA with Tukey’s multiple comparison tests was used to analyze data in which there were more than two groups. Significance was reported at *p* <0.05.

## RESULTS

### Rapid development of adaptive and innate immune suppression in sepsis

Septic mice were euthanized and spleens were harvested at six and 24 hours after CLP. Splenocytes were plated in duplicate and stimulated for 20-22 hours with either anti-CD3/CD28 mAb or LPS for evaluation of T cell and monocyte/macrophage function, respectively (**Fig.1**). Each spot represents an individual lymphocyte or monocyte/macrophage producing IFN-γ or TNF-α respectively. Sepsis caused a dramatic decrease in both the number of IFN-γ producing lymphocytes and TNF-α producing monocyte/macrophages at both time points. In addition to the decreased number of cytokine producing splenocytes, there was also a decrease in the average size of each spot indicating that the amount of IFN-γ or TNF-α produced on a per-cell basis was also reduced in septic versus naive control mice.

### Early immune suppression progresses to an increased host immune response

Additional mice were euthanized serially (days 1, 2, 3, 4, and 7 post CLP), splenocytes harvested, and IFN-γ and TNF-α production was quantitated by both ELISpot and ELISA immuno-assays (**Fig. 2**). The IFN-γ ELISpot showed a decrease in the number of lymphocytes producing IFN-γ at 24 hours after CLP but a progressive increase in IFN-γ producing lymphocytes from days 2 to 7. The TNF-α ELISpot showed a similar pattern of a progressive increase in the number of monocyte/macrophages producing TNF-α.

**Figure 2.**
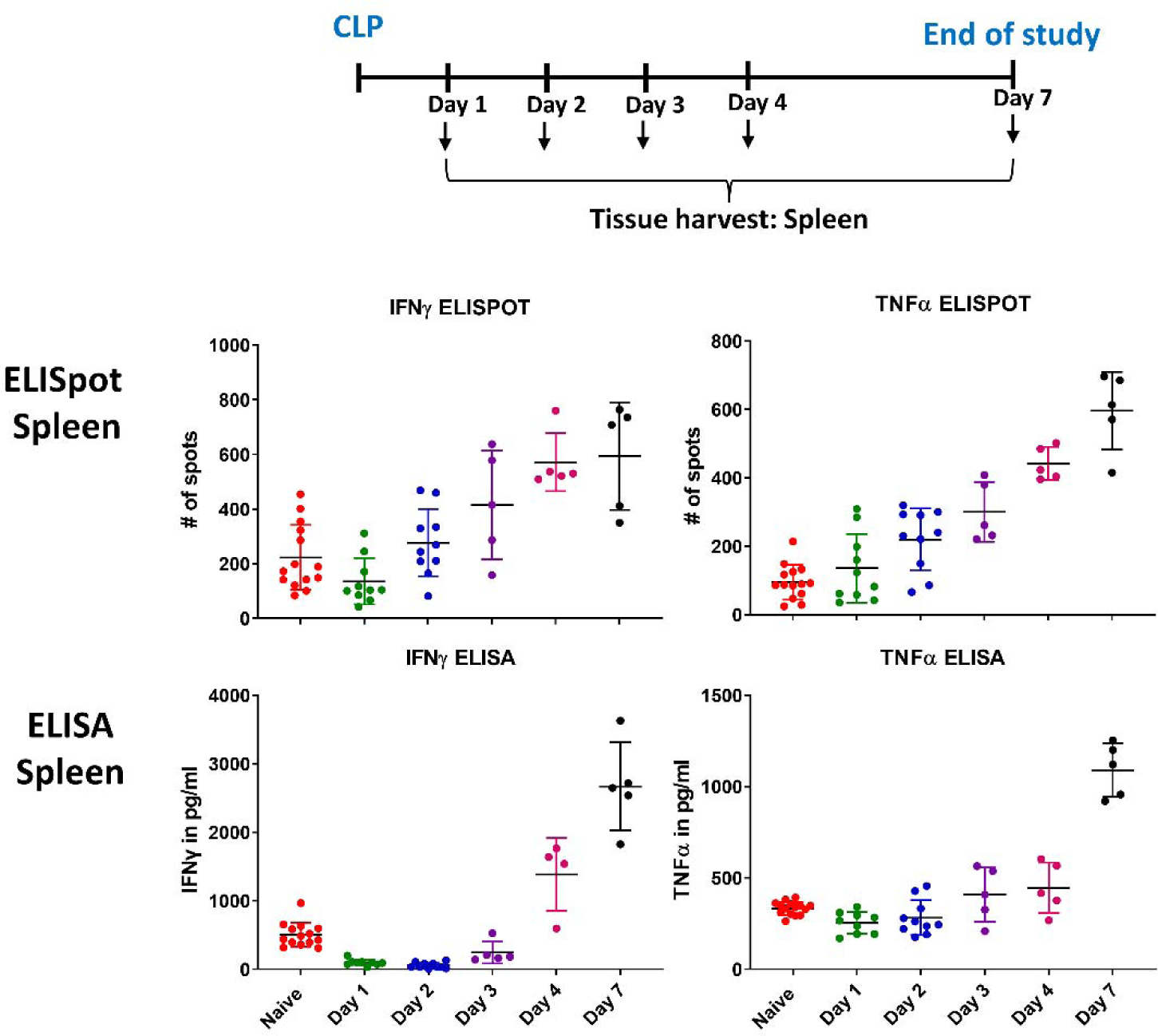
Time course for effects of sepsis on innate and adaptive immunity measured by both ELISpot and ELISA. Mice underwent CLP and spleens were harvested at days 1, 2, 3, 4, and 7 (see timeline). IFN-γ and TNF-α production from stimulated splenocytes was quantitated by both ELISpot and ELISA. Each dot represents an individual mouse. The results are the cumulative findings from three experiments.

The IFN-γ and TNF-α results obtained by the ELISpot immuno-assay were compared to the findings obtained by using the ELISA immuno-assay. Results of the IFN-γ ELISA assay showed a more sustained and deeper suppression of stimulated IFN-γ production that persisted through day 3 (**Fig. 2**). By days 4 and 7 the ELISA results for stimulated IFN-γ paralleled the findings of the ELISpot IFN-γ assay and showed a marked increase in cytokine production in septic animals versus naive controls. Of note, these immunologic results parallel the clinical condition of the septic mouse (24,25). Most septic mice in the low mortality (LD_10)_ CLP model started to recover by day four and were eating, and more active at this time point (data not shown). The results of the ELISA assay for TNF-α were different than those obtained by the ELISpot assay. The ELISA assay showed that the increase in splenocyte TNF-α production, compared to TNF-α production in naïve splenocytes, did not occur until day 7 whereas the ELISpot results showed an increased TNF-α production starting on day 2. There is a trend toward an increase in IFN-γ and TNF-α production occurring by day 4 for both ELISpot and ELISA.

### Sepsis induces a sustained loss of lymphocytes but not macrophages or dendritic cells

Flow cytometric analysis was performed on splenocytes to define the impact of sepsis on lymphocytes and monocyte/macrophage numbers that are major producers of IFN-γ and TNF-α, respectively. There was a marked decrease in CD3, CD4, and CD8 positive T cells that, although showing a trend toward recovery, tended to persist out to day 7 (**Fig. 3**). A similar significant decrease in the absolute number of splenic B cells was observed. Splenic dendritic cells were initially decreased during the first few days of sepsis but, by day 7 the number of dendritic cells had increased to a level greater than at baseline (**Fig. 3**). In contrast to the decrease in lymphocytes and dendritic cells, the number of splenic monocyte/macrophages did not decrease during sepsis and markedly increased by day 7 (**Fig. 3**).

**Figure 3.**
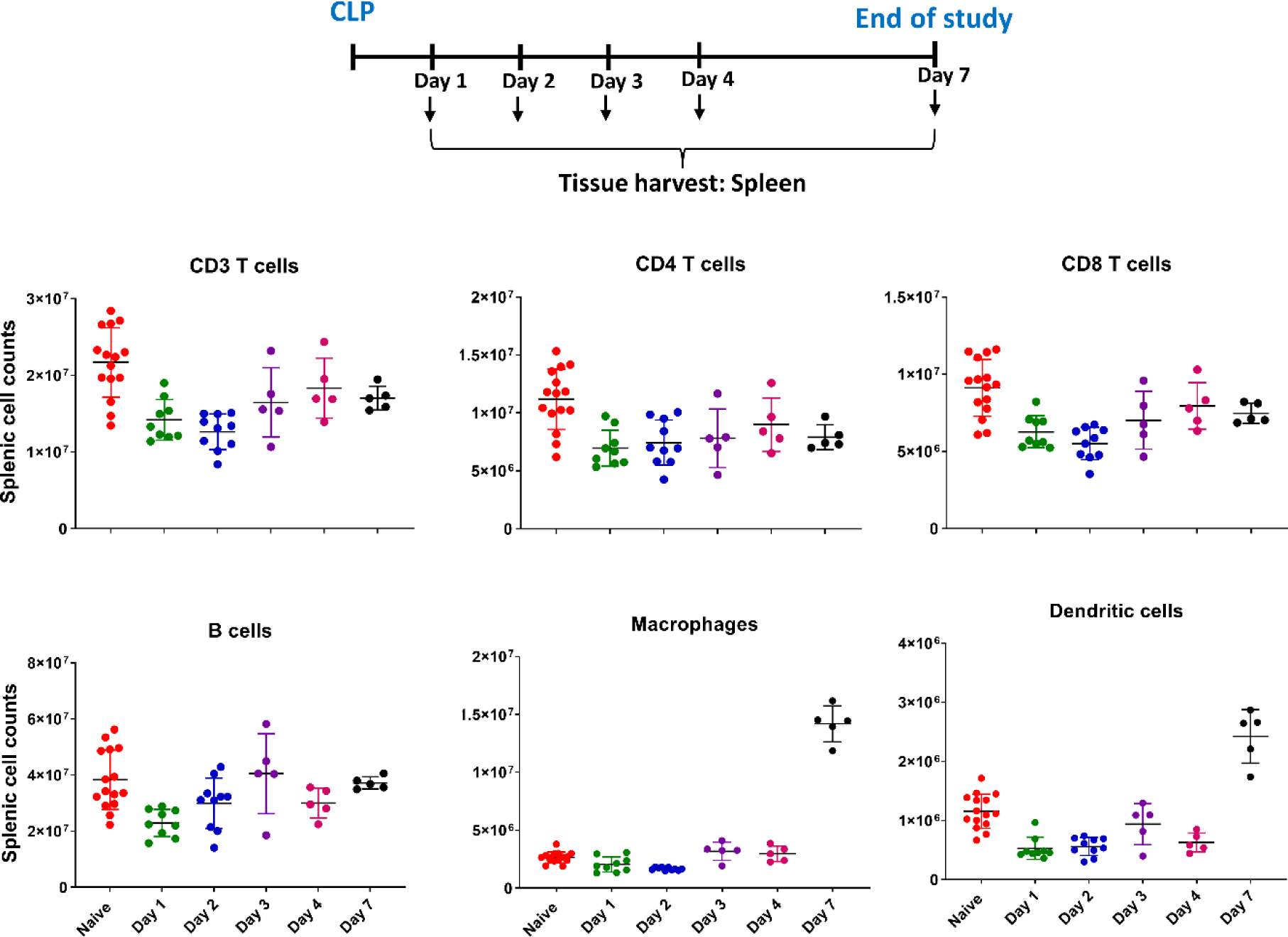
Time course for sepsis-induced loss of splenic cells. The effect of sepsis to cause loss of splenic cells was quantitated serially on days 1, 2, 3, 4, and 7. Sepsis caused a loss in absolute numbers of CD4^+^ and CD8^+^ T cells as well as B cells. The sepsis-induced loss tended to persist for up to seven days. Sepsis did not cause a loss of splenic macrophages but, in contrast, there was a remarkable increase by day 7 after CLP. There was a decrease in splenic dendritic cells that, similar to splenic macrophages, increased dramatically by day 7.

### Whole blood ELISpot showed suppressed adaptive but enhanced innate immunity

To determine whether the functionality of immune cells in the peripheral blood would reflect that of the cells in the spleen, ELISpot was also performed on whole blood from animals. Sepsis caused a marked reduction in the number of IFN-γ spot forming units in whole blood on all days tested, i.e., days 1-7 relative to the values for naive mice (**Fig. 4**). There was a trend toward improvement in IFN-γ production by day 7 but it remained lower than that obtained for naive mice. In contrast to the suppression of IFN-γ, septic mice had a marked increase in TNF-α production with the most prominent increase occurring on day 7 after CLP (**Fig. 4**).

**Figure 4.**
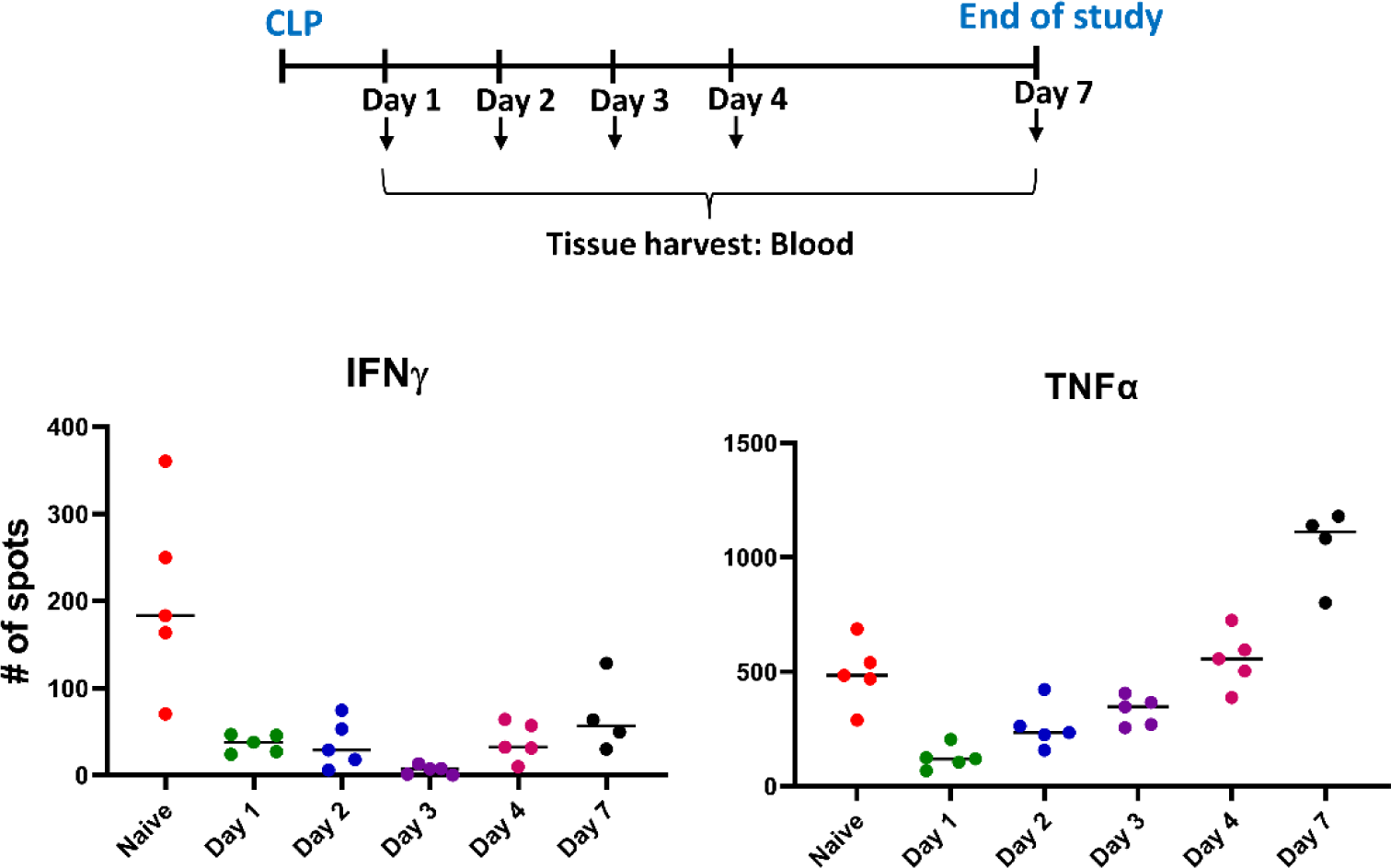
Time course for effect of sepsis on blood IFN-γ and TNF-α as detected via ELISpot. Blood was obtained serially after CLP and underwent ELISpot analysis for IFN-γ and TNF-α. Sepsis caused a profound and sustained decrease in the number of IFN-γ producing cells as well as an initial decrease in the number of TNF-α producing cells. TNF-α production recovered and exceeded the naive control group by day 7.

### Arginine and IL-7 reverse sepsis-induced immune suppression

Arginine is a metabolite essential for proper lymphocyte and monocyte/macrophage function (22). Previous work from our group showed that arginase, the enzyme that breaks down arginine, is dramatically upregulated in lymphocytes and monocytes in patients with sepsis (23). We reasoned that administration of arginine might overcome the upregulation of arginase and improve both T cell and monocyte/macrophage function in sepsis.

To examine the effect of potential immune modulatory therapies in sepsis, C57BL/6J mice underwent CLP and were treated with arginine (2.5 mg via subcutaneous injection) starting 3 hours after CLP and daily on days 1, 2, and 3 (**Fig. 5**). Splenocytes were harvested on day 5 and ELISpot assays performed. Splenocytes from septic mice treated with arginine had an increase in both IFN-γ and TNF-α production compared to septic control mice (**Fig. 5**).

**Figure. 5.**
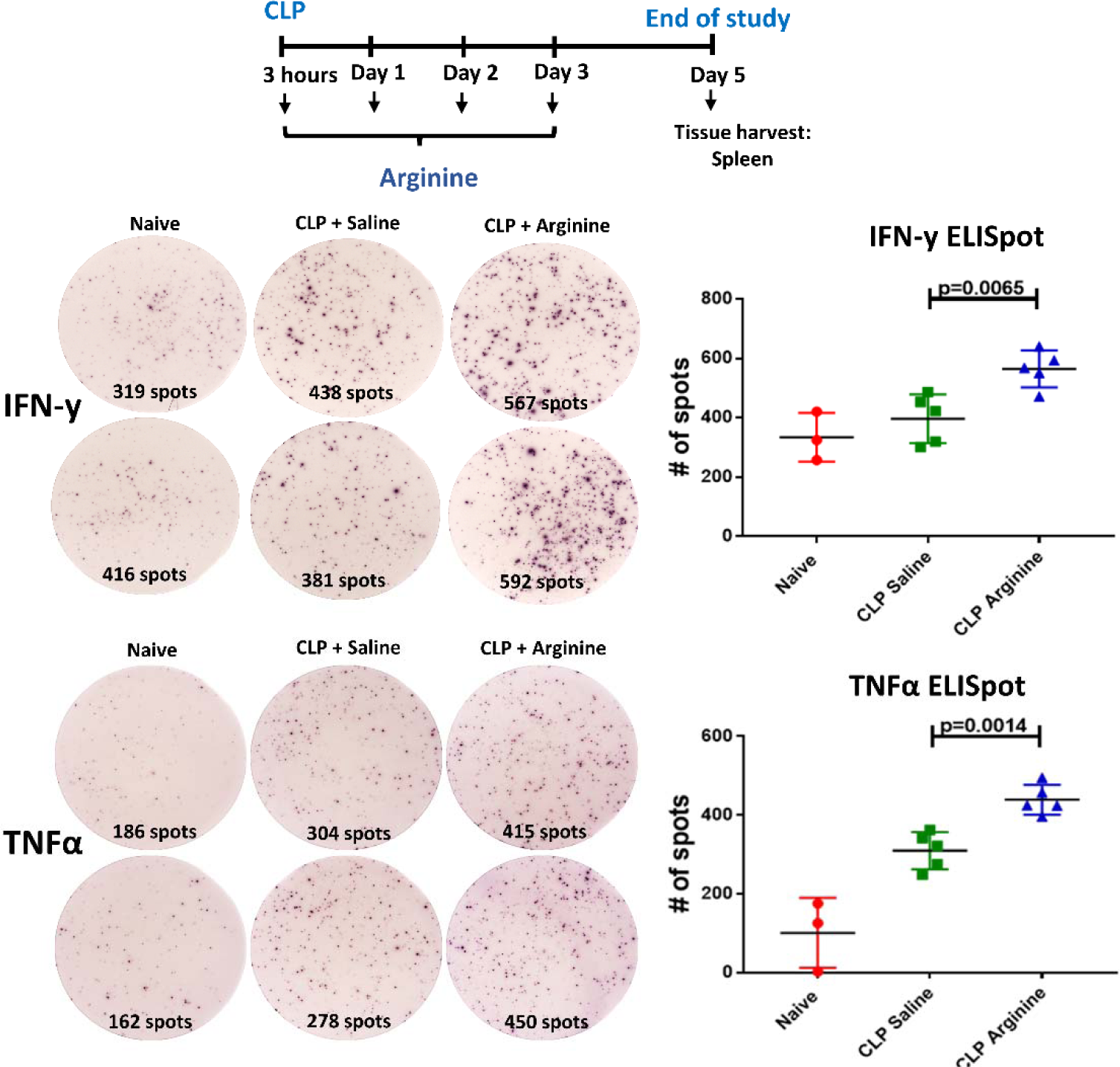
Arginine increases splenocyte IFN-γ and TNF-α production in sepsis. Mice underwent CLP and were treated with arginine at 3, 24, 48, and 72 hours after surgery. Mice treated with arginine had an increase in both IFN-γ and TNF-α production on day 5. ELISpot photomicrographs show representative images. Tests were performed in duplicate.

Additionally, ELISpot was performed on splenocytes from animals treated with the immune stimulatory agent IL-7. IL-7 is a pleiotropic cytokine that acts to increase the number and function of many subclasses of lymphocytes including CD4^+^ and CD8^+^ T cells (21). C57BL/6J mice underwent CLP and were treated with IL-7 (2.5 µg via subcutaneous injection) at 24, 28, and 72 hours after surgery (**Fig. 6**). Splenocytes were obtained on day 5 after CLP and ELISpot assay performed. Septic mice treated with IL-7 had a marked increase in the number of IFN-γ producing splenocytes versus septic mice treated with control saline diluent (**Fig. 6**). The effect of IL-7 on splenocyte production of TNF-α was not different compared to mice treated with saline diluent (data not shown).

**Figure 6.**
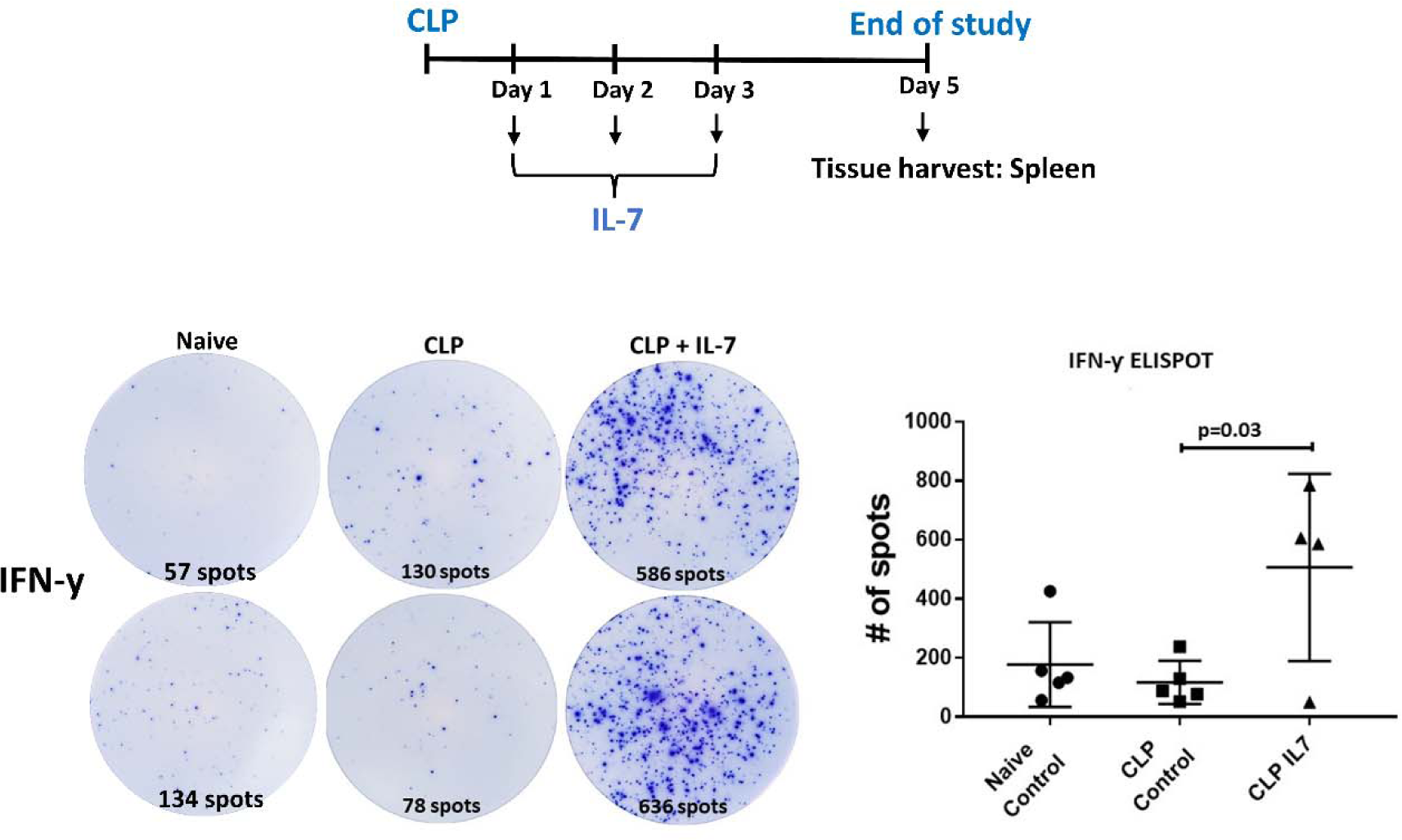
IL-7 increases splenocyte IFN-γ production in sepsis. Mice underwent CLP and were treated with IL-7 on days 1, 2, and 3. Splenocytes were obtained on day 5 and IFN-γ production assayed by ELISpot. CLP mice treated with IL-7 had increased IFN-γ production compared to mice treated with saline diluent. ELISpot photomicrographs show representative images. Tests were performed in duplicates.

### Dexamethasone induced suppression of adaptive immunity was reversed by IL-7

Dexamethasone is a corticosteroid that has potent effects to inhibit lymphocyte function and induce lymphocyte apoptosis (20). C57BL/6J mice underwent CLP and were treated with 50 μg dexamethasone in 100 μL of saline starting 24 hours after post CLP and administered daily for 4 days. We additionally tested the effect of IL-7, a potent activator of CD4 and CD8 T cells, to reverse the putative effect of dexamethasone to suppress lymphocyte function. Blood ELISpots from septic mice treated with dexamethasone showed a trend toward decreased IFN-γ production but no reduction in splenocyte IFN-γ production (**Fig. 7**). Mice treated with dexamethasone plus IL-7 showed increased production of IFN-γ in both blood and spleen (**Figs. 7**). Dexamethasone had less effects on blood and spleen ELISpot TNF-α production (**Fig. 7**). Dexamethasone can induce loss of significant numbers of lymphocytes by inducing apoptotic cell death (20). To investigate the loss of lymphocytes and its potential effect on splenocyte IFN-γ production, we analyzed splenic lymphocyte subsets from the various mouse treatment groups by flow cytometry (**Fig. 8**). Sepsis caused a trend toward decreased CD4^+^ and CD8^+^ T cell counts in the spleen which was further exacerbated by therapy with dexamethasone. IL-7 reversed the effect of dexamethasone to reduce CD8^+^ T cell counts but not splenic CD4^+^ or B cell counts (**Fig. 8**).

**Figure 7.**
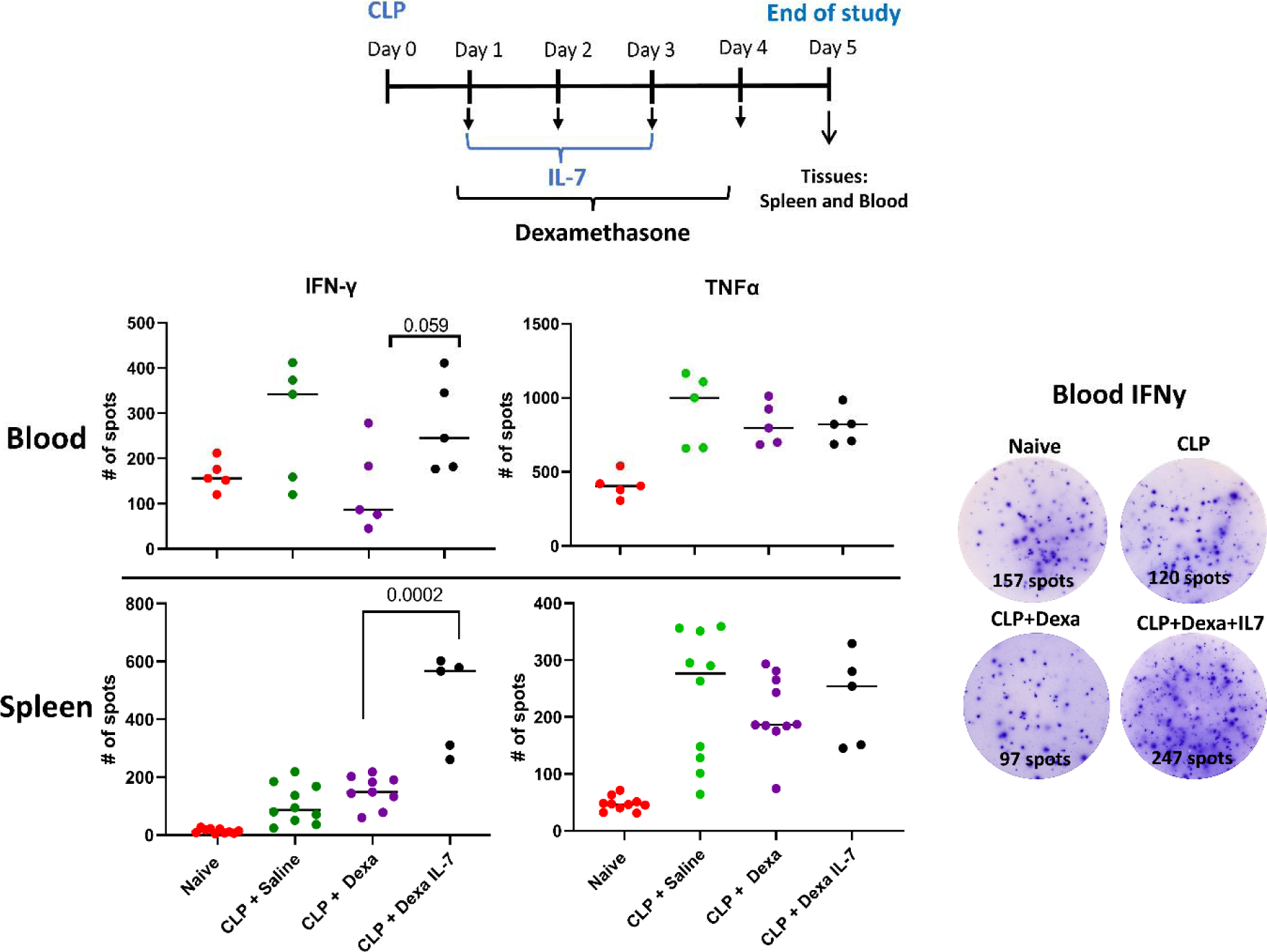
Dexamethasone and IL-7 effects on blood and spleen ELISpot IFN-γ and TNF-α production. Mice underwent CLP and were treated with dexamethasone alone or in combination with IL-7 (see timeline). Results show a trend toward decreased numbers of IFN-γ producing immune cells in blood but not spleen of septic mice in response to dexamethasone injections. IL-7 treatment combatted the effects of the dexamethasone and significantly increased the number of IFN-γ producing immune cells in spleen and demonstrated a clear trend toward increased IFN-y production in blood. Representative color microphotographs of ELISpot wells showing effects of dexamethasone and IL-7 are depicted on the right hand of the figure.

**Figure 8.**
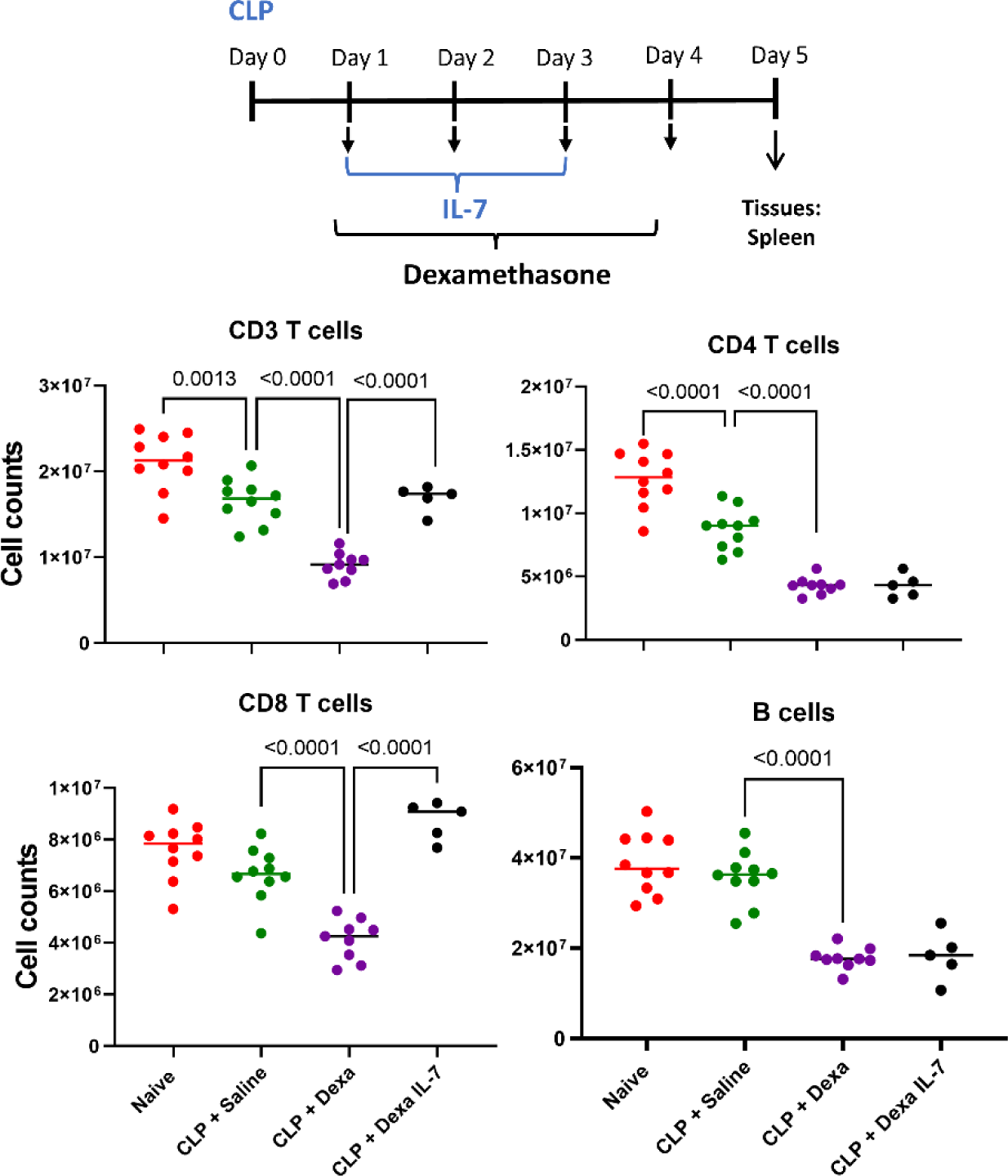
Effect of dexamethasone and IL-7 on splenic cell counts. Mice underwent CLP and were treated with dexamethasone alone or in combination with IL-7 (see timeline). Sepsis caused a loss in splenic CD3, CD4, and CD8 T cells which was markedly worsened by dexamethasone treatment. Therapy with IL-7 reversed the dexamethasone-induced CD8^+^ T cell loss but did not increase CD4^+^ T cell or B cell numbers within 5 days.

## DISCUSSION

Immunotherapy offers a new way forward in the treatment of patients with sepsis and, despite a failure of past trials to show efficacy, studies employing new immune therapies are either underway or about to begin (26,27). A key roadblock to immune therapy in sepsis in the past has been the unpredictability of the patients’ immune status and the inability to determine the immunologic endotype of patients with sepsis. Some patients with sepsis will have an over-exuberant hyper-inflammatory response and might benefit from therapy with corticosteroids or drugs which *down modulate* the immune response (2,28). Conversely, many patients will either present in an immunosuppressive state (e.g., elderly patients with immunosenescence), or develop impaired immunity as sepsis persists. These patients might benefit from immune adjuvant therapy to *enhance* their immune responsiveness. The potential importance of developing a method to determine the functional immune state of the septic patient in an actionable manner is underscored by trials of corticosteroids which show that some pediatric patients with severe sepsis/septic shock may be harmed by corticosteroids while others appear to benefit (5,6). Bline et al. used an LPS-stimulated whole blood TNF-α release assay to determine if the patients were in an immune suppressed or non-immune suppressed state (6). Those pediatric patients who had decreased TNF-α release and received corticosteroids had a longer duration of multiple organ dysfunction syndrome compared to immune suppressed patients who were not treated with corticosteroids; patients without decreased TNF-α release did not have increased risk with corticosteroid treatment (6). Based upon these and other results, the pediatric PRECISE trial intends to enroll over 1,000 pediatric patients with sepsis and employ the LPS-stimulated TNF-α response to determine if patients are immune suppressed and candidates for immune adjuvant therapy with GM-CSF (29).

There is currently an intense effort underway to develop methods to immune phenotype patients with sepsis (30,31). Many of these methods (e.g., transcriptomics, proteomics, single cell RNA sequencing), will likely be useful in predicting patient outcomes, determining molecular mechanisms of immune suppression, and in guiding new therapeutic approaches to sepsis. Most of the methods being developed, however, do not assess the actual *functional* state of the septic patient’s immune status. Knowledge of the actual functional state of the septic patient’s immunity will be essential in selecting which immune modulatory drug to use therapeutically. In fact, functional testing of the septic patient’s immune status is becoming more essential to guide immune therapies (7,8,10).

Several functional assays have been used and have identified patients who have an immunosuppressive endotype as evidenced by a decrease in stimulated production of IFN-γ or TNF-α (8,10,32). Greco et al reported that septic patients who died had a highly reduced production of IFN-γ compared to septic survivors (32). Muzhda and colleagues recently reported the results of a fully automated IFN-γ release assay in patients with sepsis (8). Septic patients had a significant decrease in IFN-γ release capacity and this finding correlated with other evidence of immune suppression including decreased monocyte HLA-DR expression.

Our group has been examining the ability of the ELISpot assay to determine the immune phenotype of patients with sepsis, i.e., whether they are in a hyper- or hypo-inflammatory phase of sepsis. The ELISpot assay has several advantages as a test of functional immunity. It has excellent dynamic range with the capability of detecting as low as a few cytokine secreting cells to over several thousand cytokine secreting cells in a five μL blood sample. Previously, we demonstrated that septic patients who had suppressed stimulated IFN-γ production had a higher mortality compared to septic patients who had a more robust stimulated IFN-γ production (10). Importantly, the ELISpot results also demonstrated the *in vitro* addition of IL-7 to the septic patient blood samples markedly improved IFN-γ production. More recently, we tested the ELISpot assay in 107 septic, 68 non-septic patients, and 46 healthy controls subjects for its ability to predict clinical outcomes (*manuscript in press Journal of Clinical Investigation Insight*). In septic patients who did not survive 180 days, stimulated whole blood IFN-γ expression was significantly reduced on ICU days 1, 4 and 7 (all *p*<0.05), due to both significant reductions in total number of IFN-γ-producing cells and in the amount of IFN-γ produced per cell (all *p*<0.05). Importantly, IFN-γ total expression on day 1 and 4 after admission could discriminate 180-day mortality better than absolute lymphocyte count, IL-6 levels, or procalcitonin.

The present results conducted in a clinically relevant animal model of sepsis show ELISpot can detect the progression of the sepsis immune response over time from a suppressed state to a hyper-inflammatory response with excellent dynamic range. The ELISpot assay was also able to distinguish the unique and at times discordant responses of the innate and adaptive immune system to sepsis. ELISpot of blood immune cells showed that there was a sustained suppression of adaptive immunity as indicated by suppressed IFN-γ production versus a lack of suppression of innate immunity as indicated by elevated TNF-α production (**Fig. 3**). Interestingly, the ELISpot IFN-γ response in splenocytes did not parallel that in blood and showed recovery of lymphocyte function beginning around day three whereas deficiency in IFN-γ production in blood persisted through day 7 (**Fig. 2 and 4**).

Additionally, the ELISpot assay was able to determine the effect of various immune active drugs to either suppress or augment the immune response (**Figs 6-8**). Although the present results showing the effect of immune adjuvants to modulate the septic immune response were conducted in a murine sepsis model, we have previously observed in two case reports that the ELISpot assay has shown enhanced T cell IFN-γ production in septic patients who were treated on a compassionate basis with IL-7 (33,34). Collectively, these studies support the notion that the ELISpot assay will be useful as a clinical tool to immune endotype septic patients, to guide appropriate immune-based drug therapies, and to follow the efficacy of the drugs in restoring immune homeostasis.

It is important to note the differences in the results of the ELISpot versus the ELISA assay as illustrated in figure 2. These differences are likely due to the fact that ELISpot detects the number of cells producing the cytokine of interest and the relative amount produced per cell, whereas ELISA is a quantitative measure of the total cytokine production. In certain situations, there may be a large number of cytokine-producing cells detected by ELISpot, but the quantitative amount of cytokine being produced by those cells could be low. In this case, ELISA would only detect a small amount of the cytokine of interest. Alternatively, there could be a few cells producing enormous amounts of cytokine, and in this case ELISpot would detect only a small number of cells whereas ELISA could detect a high concentration of released cytokine.

Despite all the strengths identified, it is important to acknowledge a number of limitations to this study. There are many differences between the mouse and human immune system and the present results showing the temporal effect of sepsis on innate and adaptive immunity in murine sepsis may not be indicative of the clinical condition in human sepsis (12). Second, the mouse CLP model of sepsis does not reflect the protracted nature of sepsis that can occur in many sepsis survivors in which chronic critical illness can persist for weeks or months. A third limitation of our study is that we did not determine the specific immune effector cells in the septic mice defective in production of IFN-γ and TNF-α. Defining which immune cells have impaired cytokine production in sepsis will help guide potential immune drug therapies that are most likely to be effective in restoring immune homeostasis.

In conclusion, the ELISpot assay demonstrated with high fidelity the dynamic changes in innate and adaptive immunity that occur with sepsis. Because ELISpot is a functional assay, it is likely that it will find its greatest use in helping guide the optimal immune drug therapy in a particular patient at a given time and in following the patient’s therapeutic response to the drug. A key test will be to determine if the results of *in vitro* testing of immune adjuvant drugs in patient blood samples are indicative of changes that occur when the patient is treated *in vivo* with the same drug therapy. Finally, we speculate the ELISpot assay or another functional test of patient immunity will have broad clinical applicability in guiding immune therapies in many disorders other than sepsis, including patients with autoimmunity, cancer, and patients who have undergone organ transplantation.

## Acknowledgements

This work was directly supported by R35 GM-12698 and R01 GM-132364

Additional support was provided by grants T32 GM-008721 (PAE), RM1 GM-139690 (LLM, PAE), R35 GM-140806 (PAE), R35 GM-0133756 (IT), R35 GM-134880 (SB), R35 GM1-40881 (TSG), from the National Institute of General Medical Sciences, and TK6BX006192 from the Department of Veteran Affairs (TSG)

